# Prediction of Michaelis constants from structural features using deep learning

**DOI:** 10.1101/2020.12.01.405928

**Authors:** Alexander Kroll, David Heckmann, Martin J. Lercher

## Abstract

The Michaelis constant *K*_M_ describes the affinity of an enzyme for a specific substrate, and is a central parameter in studies of enzyme kinetics and cellular physiology. As measurements of *K*_M_ are often difficult and time-consuming, experimental estimates exist for only a minority of enzyme-substrate combinations even in model organisms. Here, we build and train an organism-independent model that successfully predicts *K*_M_ values for natural enzyme-substrate combinations using machine and deep learning methods. Predictions are based on a task-specific molecular fingerprint of the substrate, generated using a graph neural network, and the domain structure of the enzyme. Model predictions can be used to estimate enzyme efficiencies, to relate metabolite concentrations to cellular physiology, and to fill gaps in the parameterization of kinetic models of cellular metabolism.

## Introduction

The Michaelis constant, *K*_M_, is defined as the concentration of a substrate at which an enzyme operates at half of its maximal catalytic rate; it hence describes the affinity of an enzyme for a specific substrate. Knowledge of *K*_M_ values is crucial for a quantitative understanding of enzymatic and regulatory interactions between enzymes and metabolites: it relates the intracellular concentration of a metabolite to the rate of its consumption, linking the metabolome to cellular physiology.

As experimental measurements of *K*_M_ and *k*_cat_ are difficult and time-consuming, no experimental estimates exists for many enzymes even in model organisms. For example, in *Escherichia coli*, the biochemically best characterized organism, *in vitro* measurements exist for less than 30 % of *K*_M_ values (see Methods, “Download and processing of *K_M_* values”), and turnover numbers have been measured *in vitro* for only about 10% of the ∼2000 enzymatic reactions (Davidi et al., 2016).

*K*_M_ values, together with enzyme turnover numbers, *k*_cat_, are required for models of cellular metabolism that account for the concentrations of metabolites. The current standard approach in large-scale kinetic modeling is to estimate kinetic parameters in an optimization process (Khodayari and Maranas, 2016; Saa and Nielsen, 2017; Strutz et al., 2019). These optimizations typically attempt to estimate many more unknown parameters than they have measurements as inputs, and hence the resulting *K*_M_ and *k*_cat_ values have wide confidence ranges and show little connection to experimentally observed values (Khodayari and Maranas, 2016). Therefore, predictions of these values from artificial intelligence, even if only up to an order of magnitude, would represent a major step toward more realistic models of cellular metabolism and could drastically increase the biological understanding provided by such models.

Only few previous studies attempted to predict kinetic parameters of natural enzymatic reactions *in silico*. Heckmann, Lloyd, et al. (2018) successfully employed machine learning models to predict unknown turnover numbers for reactions in *Escherichia coli*. They found that the most important predictors of *k*_cat_ were the reaction flux catalyzed by the enzyme, estimated computationally through parsimonious flux balance analysis, and structural features of the catalytic site. While many *E. coli k*_cat_ values could be predicted successfully with this model, active site information was not available for a sizeable fraction of enzymes (Heckmann, Lloyd, et al., 2018). Moreover, neither active site information nor reaction flux estimates are broadly available beyond a small number of model organisms, preventing the generalization of this approach.

Borger, Liebermeister, and Klipp (2006) trained a linear model to predict *K*_M_ values based on other *K*_M_ measurements for the same substrate paired with different enzymes in the same organism and with the same enzymes in other organisms; they fitted an independent model for each of 8 different substrates. Yan et al. (2012) later followed a similarly focused strategy, predicting *K*_M_ values of beta-glucosidases for the substrate cellobiose based on a neural network. These two previous prediction approaches for *K*_M_ targeted individual, well-studied enzyme-substrate combinations with ample experimental *K*_M_ data for training and testing. Their strategies are thus unsuitable for less well-studied reactions and cannot be applied to genome-scale predictions.

The molecular structures of the substrate and the enzyme are crucial determinants for their mutual affinity. Thus, to formulate a general predictive model for *K*_M_, it appears necessary to use aspects of the molecular structure of the substrate as input features. A widely used approach to convert molecular structures into a multidimensional numerical feature is the construction of extended-connectivity fingerprints (ECFPs) (Rogers and Hahn, 2010). To create ECFPs, an initial identifier for every atom is calculated that contains information about the atomic number, mass, and charge as well as about the number of neighboring atoms. Afterwards, these identifiers are updated for a pre-defined number of steps by iteratively applying pre-defined functions to summarize aspects of neighbouring atoms and bonds. After the iteration process, all identifiers are used as the input of a hash function to produce a binary vector with structural information about the molecule.

Recent work has shown that superior prediction performance can be achieved through a task-specific adjustment of the parameters of the pre-defined functions that are used to update the atom identifiers. Training these with deep neural networks to produce task-specific fingerprints led to state-of-the-art performances on many biological and chemical datasets (Zhou et al., 2018; Yang et al., 2019). In contrast to conventional neural networks, these graph neural networks (GNNs) can process non-Euclidean inputs such as molecules. They are thus able to learn a task-specific fingerprint of the input molecule during their optimization, simultaneously optimizing the fingerprint and using it to predict properties of the input.

To use aspects of the enzyme structure for *K*_M_ predictions, it would be desirable to employ detailed structural and physicochemical information on the substrate binding site, as done by Heckmann *et al.* for their *k*_cat_ predictions in *E. coli* (Heckmann, Lloyd, et al., 2018). However, these sites have not been characterized for a majority of enzymes (Furnham et al., 2014). An alternative approach is to employ a coarse-grained view of enzyme structure, considering only their functional domain content as represented in the Pfam database of protein domain families (Bateman et al., 2004). While this approach is capable of delivering enzyme-specific features for all enzymes, it cannot distinguish between enzyme sequences that share the same functional domain content.

*K*_M_ values are a product of evolution under natural selection (Heckmann, Zielinski, and Palsson, 2018). Consider two homologous enzymes with identical domain structure but different natural substrates, enzyme *A* with substrate *a* and enzyme *B* with substrate *b*. We expect that *A* evolved to bind *a* better than *b*, while the opposite will be true for *B*. However, a machine learning model trained on domain structures will not be able to distinguish these situations; the model will assign the same predictions for both substrates to each of the two proteins. When no sequence-specific catalytic site information is available, it is thus important to only train and test the model on natural enzyme-substrate combinations.

Here, we present a general approach to predict *K*_M_ values based on the molecular structure of the substrate and on the presence and absence of 769 functional domains in the enzyme, using machine and deep learning models. In the final model, we use information on the functional domains together with a task-specific molecular fingerprint of the substrate as the input of a gradient boosting model (T. Chen and Guestrin, 2016). Our model reaches a coefficient of determination of *R*^2^ = 0.42 between the predicted and measured values on a test set, i.e., the model explains 42% of the variability across *K*_M_ values for different substrate-enzyme combinations.

## Results

We extracted information on *K*_M_ together with the substrate name, Enzyme Commission (EC) number, and amino acid sequence of the enzyme from the BRENDA database (Jeske et al., 2019) for wild-type enzymes. Since we are only interested in one *K*_M_ value for every substrate in combination with a group of enzymes with the same functional domain content (see below), we took the geometric mean if multiple values existed for the same combination of substrate and enzyme domain content. This resulted in a data set with 5,158 data points, which was split into training, validation, and test sets (70%/10%/20%). All *K*_M_ values were log_10_-transformed.

### Predicting *K*_M_ from extended-connectivity fingerprints

We used the python package RDKit (Landrum et al., 2006) to calculate an extended-connectivity fingerprint (ECFP) for each substrate in our dataset based on the corresponding MDL Molfile, downloaded from KEGG (Kanehisa and Goto, 2000). A Molfile lists a molecule’s atom types, atom coordinates, and bond types (Dalby et al., 1992). We then used the ECFPs as the input of a fully connected neural network (FCNN), which we trained to predict the *K*_M_ values of enzyme-substrate combinations (**Figure 1a**).

**Figure 1.**
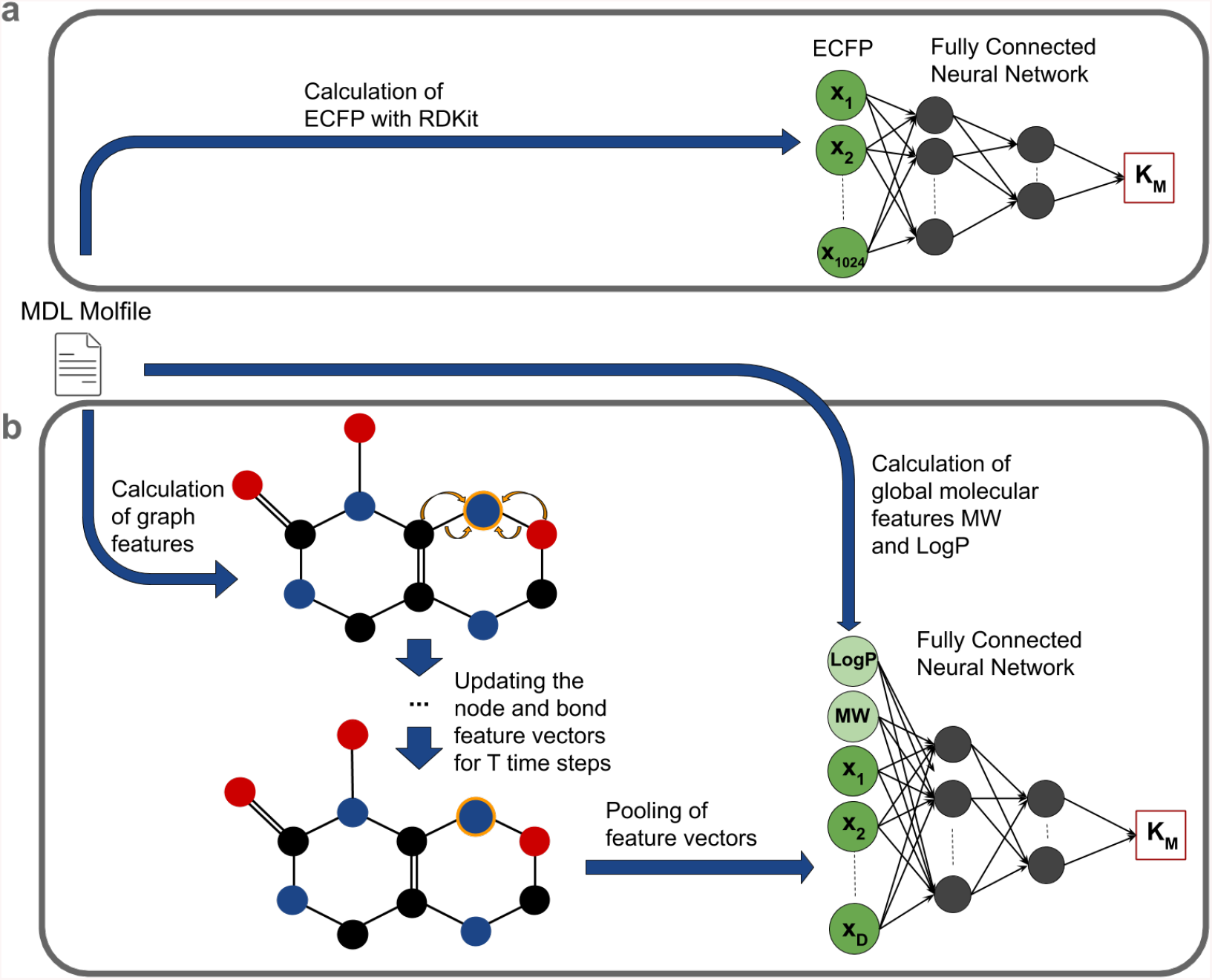
Model overview. **(a)** FCNN. 1024-dimensional ECFPs are calculated from MDL Molfiles of the substrates and then passed through the FCNN. **(b)** GNN. Node and edge feature vectors are calculated from MDL Molfiles and are then iteratively updated for *T* time steps. Afterwards, the feature vectors are pooled together into a single vector that is passed through a FCNN together with two global features of the substrate, *MW* and *LogP*.

ECFPs are binary vectors that are representations of chemical structures, calculated in an iterative process using information about a molecule’s bonds and atoms (types, masses, valences, atomic numbers, atom charges, and number of attached hydrogen atoms) (Rogers and Hahn, 2010). The number of iterations and the dimension of the fingerprint can be chosen freely. We set them to the default values of 3 and 1024, respectively; lower or higher dimensions led to inferior predictions. The FCNN consisted of a 1024-dimensional input layer, three hidden layers with dimensions 256, 64, and 32, and a 1-dimensional output layer (for more details see Methods).

The predictions based solely on ECFPs had a mean squared error *MSE* = 0.84 and a coefficient of determination *R*^2^ = 0.32 (**Figure 2**). As these predictions do not consider any enzyme-specific information, it may not be surprising that the proportion of the *K*_M_ variance explained is moderate.

**Figure 2.**
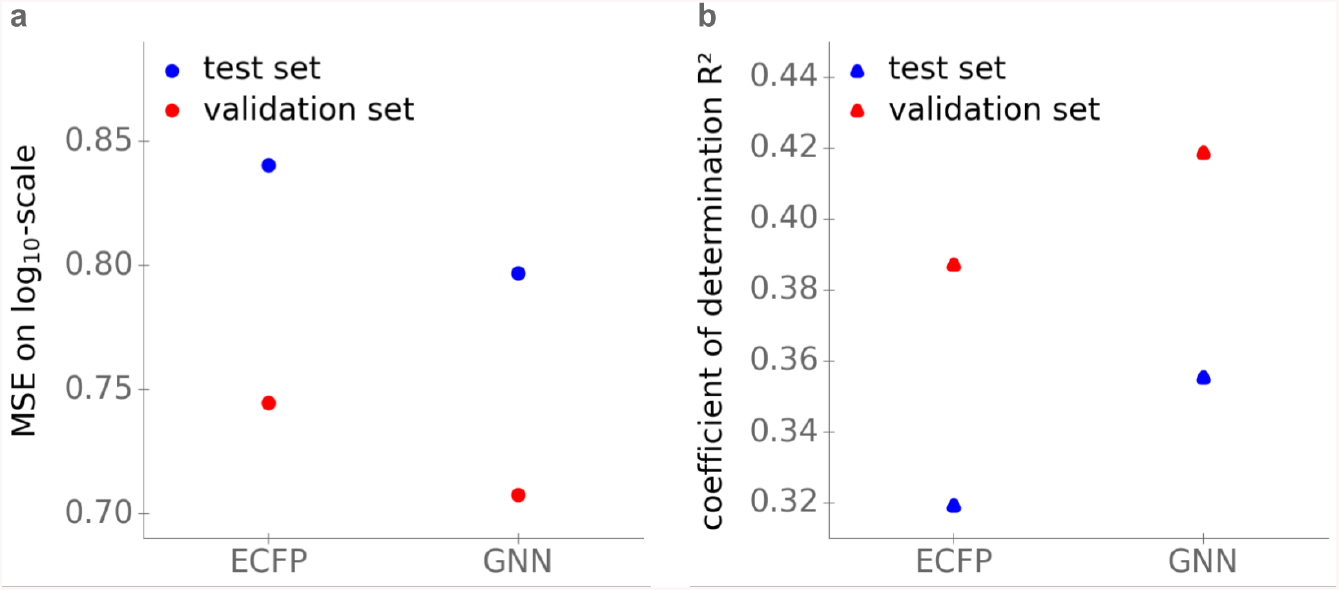
GNNs with substrates as inputs lead to superior *K*_M_ predictions compared to FCNNs with ECFPs as inputs. **(a)** Mean squared errors (*MSE*) on a log_10_-scale. **(b)** Coefficients of determination *R*^2^.

### Superior predictions using Graph Neural Networks

To test if a task-specific molecular fingerprint leads to superior *K*_M_ predictions, we next used a graph neural network (GNN) instead of the FCNN to represent the substrate molecules. A molecule is represented as a graph by interpreting the atoms as nodes and the chemical bonds as edges. We again obtained the necessary structural information for each substrate from its MDL Molfile. Before a graph is processed by a GNN, feature vectors 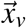 for every node *v* and feature vectors 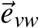 for every edge between two nodes *v* and *w* are calculated. We calculated eight features for every atom and four features for every bond of a substrate, including mass, charge, and type of atom as well as type of bond (for a full list see Methods). The initial representations 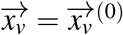 and 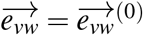 are updated iteratively for a predefined number of steps *T* using the feature vectors of the neighbouring nodes and edges (**Figure 1b**). During this process the feature vectors are multiplied with matrices with trainable entries, which are fitted during the optimization of the GNN (for more details see Methods). After *k* iterations, each node representation 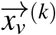 contains information about its k-hop neighbourhood graph. After completing *T* iteration steps, all node representations are averaged to obtain a single vector 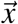, which represents the entire graph (Gilmer et al., 2017; Kearnes et al., 2016). The vector 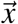 can then be used as an input of a fully connected neural network (FCNN) to predict properties of the graph (the *K*_M_ value of the molecule in our case; **Figure 1b**).

The described processing of a graph with a GNN can be divided into two phases. The first, message passing phase consists of the iteration process. The second, readout phase comprises the averaging of the node representations and the prediction of the target graph property (Gilmer et al., 2017). During the training, both phases are optimized simultaneously. The vector 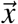 can be viewed as a task-specific fingerprint of the substrate. Since the model is trained end-to-end, the GNN learns to store all information necessary to predict *K*_M_ in this vector (Kearnes et al., 2016; Duvenaud et al., 2015).

It was shown that global molecular features can improve the performance if they are concatenated to the vector 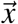 in the readout phase (Yang et al., 2019; Na, H. W. Kim, and Chang, 2020). We thus added the molecular weight (*MW*) and the octanol-water partition coefficient (*LogP*), which are global molecular features that are correlated with the *K*_M_ value (Bar-Even et al., 2011), after the message passing phase, resulting in an extended fingerprint 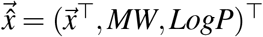. To update the feature vector representations, we performed *T* = 2 iterations using batch normalization and dropout. The dimension of *x* was chosen to be *D* = 100.

The performance of the GNN is better than that of the neural network with ECFPs, reaching a mean squared error *MSE* = 0.80 and a coefficient of determination *R*^2^ = 0.36. We verified that the difference in model performance is statistically significant, using a one-sided Wilcoxon signed-rank test for the absolute errors of the predictions for the test set (*p* = 0.0057). This shows that a task-specific molecular fingerprint contains more information about the *K*_M_ value than a predefined one. **Figure 2** compares the performances of the two trained models, GNN and FCNN, showing the mean squared errors (*MSE*) and *R*^2^ on the test set on a log_10_-scale. We note that adding the molecular weight (*MW*) and the octanol-water partition coefficient (*LogP*) to the ECFPs as extra features did not lead to improved predictions (*MSE* = 0.84, *R*^2^ = 0.32).

### Effects of molecular weight and octanol-water partition coefficient

In the calculation of the task-specific (GNN) fingerprints, we added the molecular weight (*MW*) and the water octanol-water partition coefficient (*LogP*). Do theses extra features contribute to improved *K*_M_ predictions? To answer this question, we trained GNNs without the additional features *LogP* and *MW*, as well as with only one of those additional features. **Figure 3** shows that adding the additional features has only a small effect on performance, reducing MSE from 0.82 to 0.80 while increasing *R*^2^ from 0.34 to 0.36. While the difference in model performance is statistically significant (*p* = 0.0454, one-sided Wilcoxon signed-rank test for the absolute errors of the predictions for the test set), we conclude that most of the information used to predict *K_M_* can be extracted from the graph of the molecule itself.

**Figure 3.**
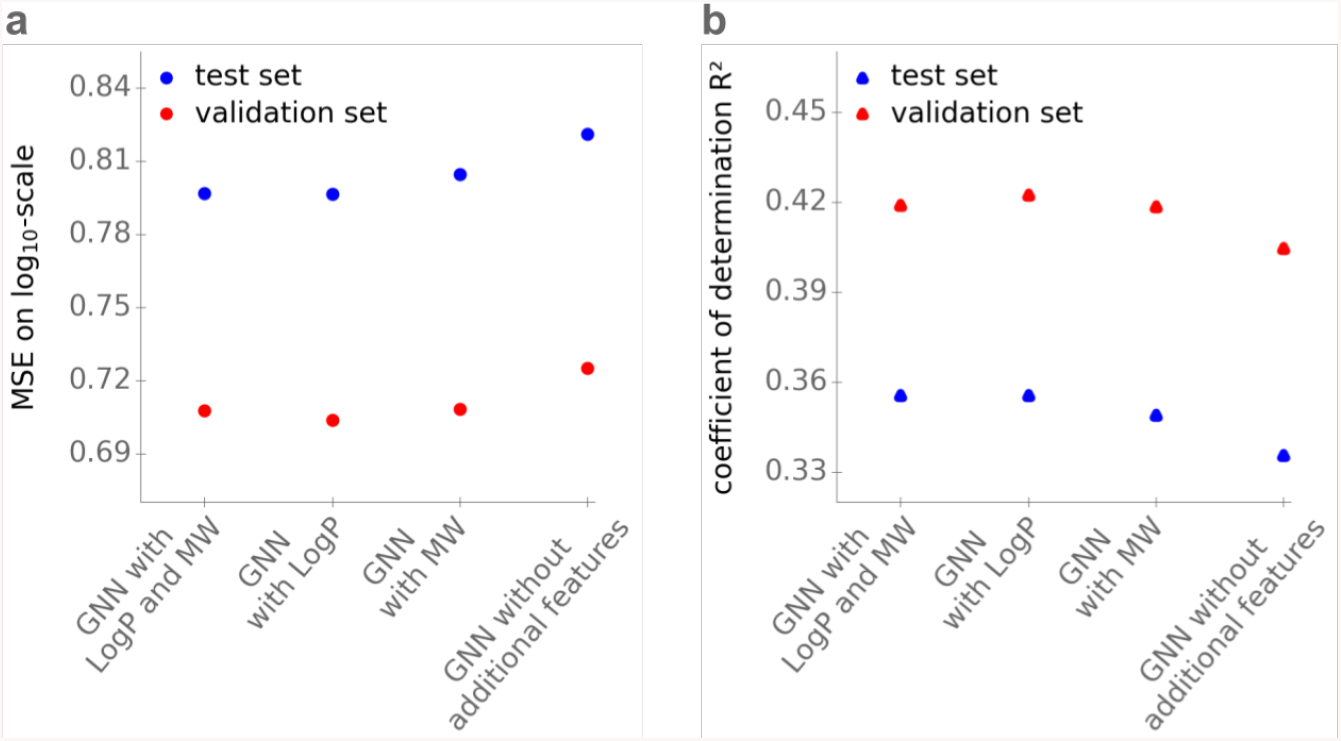
Adding the features *LogP* and *MW* has only a minor effect on the performance of the GNN in predicting *K*_M_. **(a)** Mean squared errors (*MSE*) on a log_10_-scale. **(b)** Coefficients of determination *R*^2^. Models use the GNN with additional features *LogP* and *MW*; with only one of the additional features; and without the two features.

### Functional enzyme domains as additional features

To predict *K*_M_ values for specific enzyme-substrate combinations, we need to add input features that represent enzyme properties. To avoid the drastic reduction in dataset size associated with the use of molecular features of the catalytic site, we restricted the enzyme information represented in the model to the presence of specific functional domains. To create a corresponding feature vector for each enzyme, we mapped the amino acid sequence to one or more of 18,259 functional domains from the Pfam database (Bateman et al., 2004). Initially, each enzyme was represented by an 18,259-dimensional binary vector (Y. Li et al., 2018) (for details see Methods). Disregarding all entries belonging to functional domains with less than 5 occurrences in our whole dataset reduced this to a 769-dimensional binary vector for every enzyme.

Since we only use information about the functional domains of an enzyme, we cannot distinguish between different enzymes with identical sets of domains. To improve legibility, in the following we will refer to such a group of enzymes that share the same functional domain set simply as an enzyme.

### Predicting *K*_M_ using substrate and enzyme information

To predict the *K*_M_ value, we concatenated the 102-dimensional task-specific extended fingerprint learned with the GNN and the 769-dimensional vector with information about the enzyme domain content. This feature vector was then used as the input for a gradient boost model for regression. Gradient boosting is a technique that creates an ensemble of many decision trees to make predictions. We also trained a fully connected neural network; however, predictions were substantially worse.

The gradient boost model that combines substrate and enzyme information achieves a mean squared error *MSE* = 0.72 on a log_10_-scale and results in a coefficient of determination *R*^2^ = 0.42, substantially superior to the above models based on substrate information alone. **Figure 4a** and **4b** compare the performance of the full model to models that only use substrate or enzyme information as input on a test dataset of previously unseen enzyme-substrate combinations. To predict the *K*_M_ value with only enzyme information as its input, we again fitted a gradient boost model, leading to *MSE* = 1.03 and *R*^2^ = 0.17. To predict the *K*_M_ value from substrate information only, we chose the GNN that was used to create the task-specific fingerprint of the substrate (**Figure 2**).

**Figure 4.**
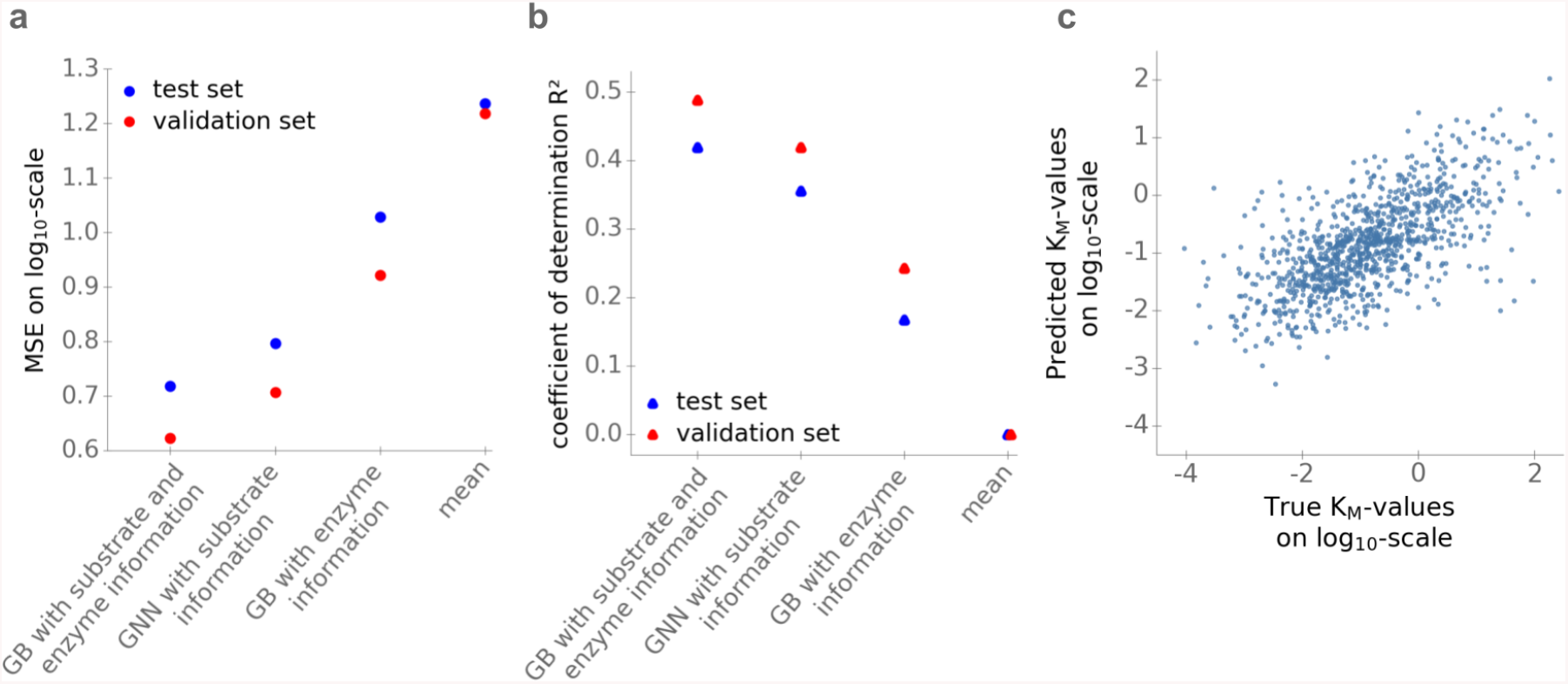
Performance of the optimized models. **(a)** Mean squared errors (*MSE*). **(b)** Coefficients of determination (*R*^2^). Values in (a) and (b) are calculated for the validation and test sets, using the gradient boost model with different inputs: substrate and enzyme information; substrate information only (GNN); enzyme information only (domain content). For comparison, we also show results from a naïve model using the mean of the *K*_M_ values in the training set for all predictions. **(c)** Scatter plot of log_10_-transformed *K*_M_ values of the test set predicted with the gradient boost model with substrate and enzyme information as inputs versus the experimental values downloaded from BRENDA.

**Figure 4a** and **4b** also compare the three models to the naïve approach of simply using the mean over all *K*_M_ values in the training set as a prediction for all *K*_M_ values in the test set, resulting in *MSE* = 1.23 and *R*^2^ = 0. **Figure 4c** compares the values predicted using the full model with the experimental values of the test set obtained from BRENDA.

### Predicting *K*_M_ for enzymes and substrates not represented in the training data

Homologous enzymes that catalyze the same reaction in different organisms tend to have broadly similar kinetic parameters. However, since the test set contains only three data points with substrate-EC number combinations that also appear in the training set (for a different organism), inclusion of such data points cannot explain our results.

It is conceivable, however, that predictions are substantially better if the training set contains entries with the same substrate or with the same enzyme, even if not in the same combination. In practice, one may want to also make predictions in cases where the enzyme and/or the substrate are not represented in the training data. To test how our model performs in such cases, we separately analysed those *n* = 9 entries in the test data where neither enzyme nor substrate occurred in the the training data, resulting in *MSE* = 0.53 and *R*^2^ = 0.35, compared to *MSE* = 0.72 and *R*^2^ = 0.42 for the full test data. To enlarge the sample size for this test, we created additional training and test sets, where the test set consists of *n* = 45 enzyme-substrate combinations completely disjunct to the substrates and enzymes in the training set (for details see Methods). Training a model on the new training set led to *MSE* = 0.73 and *R*^2^ = 0.32 on the disjunct test set. We thus conclude that the proposed model appears capable of predicting *K*_M_ values independent of the occurrence of enzyme and/or substrate in the training data.

## Discussion

In conclusion, we found that Michaelis constants *K*_M_ of enzyme-substrate pairs can be predicted with satisfactory accuracy through machine learning, reaching *R*^2^ = 0.42 between predicted and measured values on a test dataset not used for model training. To obtain this predictive performance, we used task-specific fingerprints of the substrate (GNN) optimized for the *K*_M_ prediction, as these apparently contain more information about *K*_M_ values than predefined molecular fingerprints based on expert-crafted transformations (ECFP). The superior performance of GNNs is in line with the results of a previous study, which compared the performance of GNNs and predefined molecular fingerprints on the prediction of chemical characteristics of small molecules (Yang et al., 2019).

A direct comparison to the results of Yan et al. (2012) is not possible, as they trained reaction-specific models that specifically aim to distinguish *K*_M_ values between different sequences of the same enzyme. However, the performance of our general model, with mean squared error *MSE* = 0.72, compares favorably to that of the substrate-specific models of Borger, Liebermeister, and Klipp (2006), which resulted in an overall *MSE* = 1.02.

The only information about the enzyme that is used in our model is the presence and absence of functional domains. It appears somewhat surprising that this simple feature set provides important enzyme-specific information for the prediction of *K*_M_. Obviously, *K*_M_ values can differ between homologous enzyme sequences with identical domain structures, e.g., for alleles or paralogs within one species or for orthologs between species. Predictions could hence be improved by taking into account allele-specific information, such as the hydrophobicity, depth, or structural features (Heckmann, Lloyd, et al., 2018) of the enzyme active site, once such features become widely available (Furnham et al., 2014). Models might be further improved by adding organism-specific information, such as the typical intracellular pH or temperature.

To put the performance of the current model into perspective, we consider the mean relative prediction error *MRPE* = 4.3, meaning that our predictions deviate from experimental estimates on average by 4.3-fold. This compares to a mean relative deviation of 3.7-fold between a single *K*_M_ measurement and the mean of all other measurements for the same enzyme-substrate combination in the BRENDA dataset; the latter mean is what was used for training the models. While part of the high variability across values in BRENDA is due to differences between homologous enzyme sequences, varying assay conditions in the *in vitro* experiments also contribute substantially to the variation (Bar-Even et al., 2011). Moreover, entries in BRENDA are not free from errors; on the order of 10% of the values in the database do not correspond to the values in the original papers, e.g., due to errors in unit conversion (Bar-Even et al., 2011).

Especially on the background of this variation, the performance of our enzyme-substrate specific *K*_M_ model appears remarkable. In contrast to previous attempts, the model requires no previous knowledge about measured *K*_M_ values for the considered substrate or enzyme. Furthermore, only one general-purpose model is trained, and it is not necessary to obtain training data and to fit new models for individual substrates or enzyme groups. Once the model has been fitted, it can provide genome-scale *K*_M_ predictions from existing features within minutes.

Any quantitative description of enzymatic reaction systems that explicitly considers reactant concentrations requires reliable estimates of *K*_M_ values. Predictions with the model proposed here could replace ad-hoc kinetic parameterizations, or could serve as initial values and to create boundaries for parameter estimation procedures. Future generalizations of our approach that consider additional enzyme sequence-specific features would be able to distinguish between homologous enzymes with the same domain architecture, and could be used to identify candidate substrates for uncharacterized enzymes or side reactivities of known enzymes.

## Methods

### Software and code availability

We implemented all code in Python (Van Rossum and Drake, 2009). We implemented the neural networks using the deep learning library TensorFlow (M. Abadi et al., 2015) and Keras (Chollet et al., 2015). We fitted the gradient boost models using the library XGBoost (T. Chen and Guestrin, 2016).

The Python code used to generate the results is available in the form of four Jupyter notebooks, available from https://github.com/AlexanderKroll/KM_prediction. One of these note-books contains all the necessary steps to download the data from BRENDA and to preprocess it. Execution of a second notebook performs training and validation of our final model. Two additional notebooks contain code to train the fully-connected neural network (FCNN) with ECFPs as inputs and to investigate the effect of the two additional features, *MW* and *LogP*, for the GNN.

### Download and processing of *K*_M_ values

We downloaded *K*_M_ values together with the substrate name, EC number, and amino acid sequence of the enzyme from BRENDA (Jeske et al., 2019). This resulted in a dataset with 130,456 entries. We mapped substrate names to KEGG Compound IDs via a synonym list from KEGG (Kanehisa and Goto, 2000). For all substrate names, that could not be mapped to a KEGG Compound ID directly, we tried to map them first to PubChem Compound IDs via a synonym list from PubChem (S. Kim et al., 2019) and then mapped these IDs to KEGG Compound IDs using the webservice of MBROLE (López-Ibáñez, Pazos, and Chagoyen, 2016). Then we removed: (i) all duplicates (i.e., entries with identical values for *K*_M_, substrate, EC number, and organism as another entry); (ii) all entries with non-wild type enzymes (i.e., with a commentary field in BRENDA labeling it as mutant or recombinant); (iii) entries without an amino acid sequence in BRENDA; and (iv) entries with substrate names that could not be mapped to a KEGG Compound ID. This resulted in a filtered set of 57,744 data points. For 27,032 of these, we could find an entry for the EC number-substrate combination in the KEGG reaction database. Since we are only interested in *K*_M_ values for natural substrates, we only kept these data points (Bar-Even et al., 2011). We log_10_-transformed all *K*_M_ values in this dataset.

The most detailed of our machine learning models uses as input features the molecular fingerprint of the substrate and the functional domain content of the enzyme. Thus, none of our models can distinguish between enzyme sequences with identical domain contents. If multiple log_10_-transformed *K*_M_ values existed for one substrate and one domain content (see below), we took the mean across these values, which resulted in a final dataset with 5,158 data points. We split the dataset randomly in training data (70%), validation data (10%) and test data (20%). We used the training data to fit the parameters of the machine learning models, the validation data to evaluate the models during hyperparameter optimization, and the test data to evaluate the final models after hyperparameter optimization.

To estimate the proportion of metabolic enzymes with *K*_M_ values measured *in vitro* for *E. coli*, we mapped the *E. coli K*_M_ values downloaded from BRENDA to reactions of the genome scale metabolic model *i*ML1515 (Monk et al., 2017), which comprises over 2,700 different reactions. To do this, we extracted all enzyme-substrate combinations from the *i*ML1515 model for which the model annotations listed an EC number for the enzyme and a KEGG Compound ID for the substrate, resulting in 2,656 enzyme-substrate combinations. For 795 of these combinations (i.e. 29,93 %), we were able to find a *K*_M_ value in the BRENDA database.

### Calculation of extended-connectivity fingerprints

We first representend each substrate through its extended-connectivity fingerprint (ECFP). For every substrate in the final dataset, we downloaded an MDL Molfile with two-dimensional projections of its atoms and bonds from KEGG (Kanehisa and Goto, 2000) via the KEGG Compound ID. We then used the package Chem from RDKit (Landrum et al., 2006) to calculate the 1024-dimensional binary ECFP (Rogers and Hahn, 2010) with a radius of 3 and the Molfile as the input.

### Architecture of the fully connected neural network with ECFPs

We used a fully connected neural network (FCNN) to predict *K*_M_ values using only the 1024-dimensional ECFPs as input features. The FCNN consisted of three hidden layers with dimensions 256, 64 and 32. We used rectified linear units (*ReLUs*), which are defined as *ReLU* (*x*) = *max*(*x,* 0), as activation functions in the hidden layers to introduce non-linearity. We applied batch normalization (Ioffe and Szegedy, 2015) after each hidden layer. Additionally, we used *L*2-regularization in every layer to prevent overfitting. Adding dropout (Srivastava et al., 2014) did not improve the model performance. We optimized the model by minimizing the mean squared error (*MSE*) with the Stochastic Gradient Descent with Nesterov Momentum as an optimizer with a learning rate of 10^−4^ and a momentum of 0.8. We trained the model for 50 epochs with a batch size of 32 and with a regularization parameter of *λ* = 2. The hyperparameters regularization factor, learning rate, dropout rate, batch size, and momentum were optimized by performing a grid search. We selected the set of hyperparameters with the lowest *MSE* on the validation set.

### Calculation of molecular weight (*MW*) and the octanol-water partition coefficient (*LogP*)

We calculated the additional two molecular features, molecular weight (*MW*) and the octanol-water partition coefficient (*LogP*), with the package Chem from RDKit (Landrum et al., 2006), with the MDL Molfile of the substrate as the input.

### Calculation of the input of the Graph Neural Network

Graphs in GNNs are represented with tensors and matrices. To calculate the input matrices and tensors, we used the package Chem from RDKit (Landrum et al., 2006) with MDL Molfiles of the substrates as inputs to calculate eight features for very atom *v* (atomic number, number of bonds, charge, number of hydrogen bonds, mass, aromaticity, hybridization type, chirality) and four features for every bond between two atoms *v* and *w* (bond type, part of ring, stereo configuration, aromaticity). Converting these features (except for atom mass) into one-hot encoded vectors resulted in a feature vector with *F_b_* = 10 dimensions for every bond and in a feature vector with *F_a_* = 32 dimensions for every atom.

For a substrate with *N* atoms, we stored all bonds in an *N* × *N*-dimensional adjacency matrix *A*, i.e. entry *A_vw_* is equal to one if there is a bond between the two atoms *v* and *w* and zero otherwise. We stored the bond features in a (*N* × *N* × *F_b_*)-dimensional tensor *E*, where entry 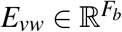 contains the feature vector of the bond between atom *v* and atom *w*. Afterwards, we expanded tensor *E* by concatenating the feature vector of atom *v* to the feature vector *E_vw_*. If there was no bond between the atoms *v* and *w*, i.e. *A_vw_* = 0, we set all entries of *E_vw_* to zero. We then used the resulting (*N* × *N* × (*F_a_* + *F_b_*))-dimensional tensor *E*, together with the adjacency matrix *A*, as the input of the GNN.

During training, the number of atoms *N* in a graph has to be restricted to a maximum. We set the maximum to 70, which allowed us to include most of the substrates in the training. After training, the GNN can process substrates of arbitrary sizes.

### Architecture of the Graph Neural Network

We use a variant of GNNs called Directed Message Passing Neural Network (D-MPNN) (H. Dai, B. Dai, and Song, 2016; Yang et al., 2019). In D-MPNNs, every edge is viewed as two directed edges pointing in opposite directions. During the iteration process (the message passing phase), feature vectors of nodes and edges are iteratively updated. To update them, feature vectors of neighbouring nodes and edges are multiplied by matrices with learnable parameters and the results are summed. Then, an activation function, the rectified linear unit (*ReLU*), is applied to the resulting vector to introduce non-linearities.

We set the number of iterations for updating the feature vector representations to *T* = 2. The dimension of the feature vectors during the message passing phase was set to *D* = 100. We applied batch normalization before every activation function. Additionally, dropout (with a droprate of 0.2) was applied at the end of the message passing phase.

After the message passing phase, the readout phase starts, and feature vectors of all nodes and edges are pooled together using an order-invariant function to obtain a single vector 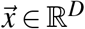, which is a representation of the input. The pooling is done using the element-wise mean of the feature vectors.

We then concatenate 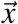 with the molecular weight (*MW*) and the octanol-water partition coefficient (*LogP*), which are global molecular features that are correlated with the *K*_M_ value (Bar-Even et al., 2011). This results in an extended fingerprint 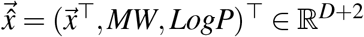.

Afterwards 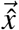 is used as the input of a FCNN with two layers with dimensions 32 and 8, again using *ReLUs* as activation functions. Batch normalization and L2-regularization are applied to the fully connected layers to avoid overfitting.

During training, the values of the matrices from the message passing phase and the parameters of the FCNN from the readout phase are fitted simultaneously. We trained the model by minimizing the mean squared error (*MSE*) with the optimizer Adadelta (Zeiler, 2012) with a decaying learning rate (decay rate to *ρ* = 0.95), with two subsequent rounds starting at 0.1 for 40 epochs and then starting at 0.01 for 10 epochs. We used a batch size of 64 and a regularization parameter *λ* = 0.2. The hyperparameters regularization factor, learning rate, batch size, dimension of feature vectors *D*, dropout rate, and decay rate were optimized by performing a grid search. We selected the set of hyperparameters with the lowest *MSE* on the validation set.

### Functional enzyme domains

To obtain the functional domains of enzymes, we stored the amino acid sequences in files in FASTA format (Lipman and Pearson, 1985). We then used the files as an input for the webservice of Pfam (Bateman et al., 2004). Pfam uses the HMMER website (Eddy, 1998) to map each amino acid sequence to one or more of 18,259 domains. For each enzyme sequences, we stored the presence and absence of functional domains in an 18,259-dimensional binary vector, where each entry represents one domain. Disregarding all functional domains with less than 5 occurrences in our whole dataset resulted in a final 769-dimensional binary vector 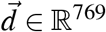 for every enzyme.

### Fitting of the gradient boost model

We concatenated the task-specific substrate fingerprint 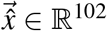 and the binary vector 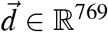 with information about the enzyme’s functional domains. We used the resulting 871-dimensional vector as the input for a gradient boost model for regression, which we trained to predict the *K*_M_ value. We fitted the model using the gradient boost library XGBoost (T. Chen and Guestrin, 2016) for Python. We set the maximal tree depth to 4, minimum child weight to 1, the learning rate to 0.04, the subsample coefficient to 0.9, and the regularization coefficient to 0.8. We trained the model for 1600 iterations. The hyperparameters regularization coefficient, learning rate, maximal tree depth, subsample coefficient, and minimum child weight were optimized by performing a grid search. We selected the set of hyperparameters with the lowest *MSE* on the validation set.

### Obtaining a test set with unseen enzymes and substrates

To test the prediction performance for enzyme-substrate combinations for which neither enzyme nor substrate appear in the training set, we created a second pair of training and test sets. We randomly chose one substrate-enzyme combination for the test set. We then also placed all data points with the same enzyme or with the same substrate in the test set. We iteratively repeated the last step for all data points newly added to the test set until no new data points were added to the test set, thereby ensuring that the training set did not contain any enzymes or substrates that also appear in the test set. For almost all of the data points this resulted in a test set containing 5, 113 out of 5, 158 data points. The maximal size of the test set that we could achieve without placing over 99% of the data points in the test set was 45; this set was chosen for further analysis.

### Model comparison

To test if the difference in performance between the GNN with substrates as input and the FCNN with ECFPs as input is statistically significant, we applied a one-sided Wilcoxon signed-rank test. We tested the null hypothesis that the median of the absolute errors on the test set for predictions made with the GNN, 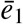, is greater or equal to the corresponding median for predictions made with the FCNN, 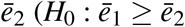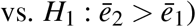. We could reject *H*_0_ (*p* = 0.0057), accepting the alternative hypothesis *H*_1_.

Analogous to the described procedure, we tested if the difference in model performance between the GNNs with and without the two additional features, *MW* and *LogP*, is statistically significant. We could reject the null hypothesis *H*_0_ that the median of the absolute errors on the test set for predictions made with the GNN with *MW* and *LogP* is greater or equal to the corresponding median for predictions made with the GNN without additional feature (*p* = 0.045)s. To execute the tests, we used the Python library SciPy (Virtanen et al., 2020).

## Acknowledgements

We thank Hugo Dourado, Markus Kollmann, and Kyra Mooren for helpful discussions. This work was funded by the Volkswagenstiftung and by the Deutsche Forschungsgemeinschaft (DFG, German Research Foundation) through grant CRC 1310 and, under Germany’s Excellence Strategy, through grant EXC 2048/1 (Project ID: 390686111).

## Author Contributions

AK performed the analyses and wrote the manuscript. AK, DH, and MJL designed the study, interpreted the results, and edited the manuscript.

